# Quantifying climatic and socio-economic influences on urban malaria in Surat, India: a modelling study

**DOI:** 10.1101/583880

**Authors:** Mauricio Santos-Vega, Rachel Lowe, Luc Anselin, Vikas Desai, Keshav G. Vaishnav, Ashish Naik, Mercedes Pascual

## Abstract

**Background:** Cities are becoming increasingly important habitats for mosquito-borne infections. The pronounced heterogeneity of urban landscapes challenges our understanding of the spatio-temporal dynamics of these diseases, and of the influence of climate and socio-economic factors at different spatial scales. Here, we quantify this joint influence on malaria risk by taking advantage of an extensive dataset in both space and time for reported *Plasmodium falciparum* cases in the city of Surat, Northwest India.

**Methods:** We analyzed 10 years of monthly falciparum cases resolved at three nested spatial resolutions (for 7 zones, 32 units and 478 workers unit’s subdivisions, respectively). With a Bayesian hierarchical mixed model that incorporates effects of population density, poverty, humidity and temperature, we investigate the main drivers of spatio-temporal malaria risk at the intermediate scale of districts. The significance of covariates and the model fit is then examined at lower and higher resolutions.

**Findings:** The spatial variation of urban malaria cases is strongly stationary in time, whereby locations exhibiting high and low yearly cases remain largely consistent across years. Local socio-economic variation can be summarized with two main principal components, representing poverty and population density respectively. The model that incorporates these two factors together with local temperature and global relative humidity, best explains monthly malaria patterns at the intermediate resolution. The effects of local temperature and population density remain significant at the finest spatial scale. We further identify the specific areas where such increased resolution improves model fit.

**Interpretation:** Malaria risk patterns within the city are largely driven by fixed spatial structures, highlighting the key role of local climate conditions and social inequality. As a result, malaria elimination efforts in the Indian subcontinent can benefit from identifying, predicting and targeting disease hotspots within cities. Spatio-temporal statistical models for the mesoscale of administrative units can inform control efforts, and be complemented with bespoke plans in the identified areas where finer scale data could be of value.

**Research in context:** *Evidence before this study:* Urban areas have become the new dominant ecosystem around the globe. Developing countries comprise the most urbanized regions of the world, with 80% of their population living in cities and an expected increase to 90% by 2050. The large and heterogeneous environments of today challenge the understanding and control of infectious disease dynamics, including of those transmitted by vectors. Malaria in the Indian subcontinent has an important urban component given the existence of a truly urban mosquito vector *Anopheles stephensi*. A literature search in Mendeley of “urban malaria” and “India” returned 161 publications, in their majority on diagnostics or brief reports on the disease, and on cross-sectional rather than longitudinal studies addressing the spatio-temporal variation of disease risk for a whole city, the subject of our work. A relevant exception is a study for the city of Ahmedabad; this not address multiple seasons across different spatial scales, and climatic conditions are not considered jointly with socio-economic drivers in the modeling. A second Mendeley search on *A. stephensi* returned 11 publications into two distinct groups: early entomological studies for India and recent reports of the mosquito in the Horn of Africa. This geographical expansion makes the specter of urban malaria a future possibility for the African continent where the disease remains so far rural and peri-urban.

*Added value of this study:* This paper relies on an extensive surveillance data set of *Plasmodium falciparum* cases for Surat (India) to investigate the variation and drivers of malaria risk in an heterogenous urban environment. A statistical model for the spatio-temporal variability of cases is developed, which includes both climatic and socio-economic drivers, with the latter summarized into two major axes of variation. Model fits are compared across three spatial resolutions, ranging from a few zones to a few hundred units. Seasonal hotspots are shown to be largely stationary in time, which allows identification of dominant drivers, including population density and local temperatures, whereas humidity acts globally modulating year-to-year burden. More granular statistical models and datasets like the one analyzed here are needed to capture the effects of socioeconomic and climatic drivers, and to predict current and future malaria incidence patterns within cities.

*Implications of all the available evidence:* The analysis identifies relevant resolution which can vary across the city for targeted intervention, including vector control, that would focus on reducing and eliminating transmission hotspots. The modeling framework, incorporating predictors representing climate at local vs. aggregate levels, and major axes of socio-economic variation, should apply to other vector-borne diseases and other cities for which surveillance records are available. The importance of spatially-explicit and sustained surveillance data for informing these models cannot be overstated.

## Background

Cities are becoming a dominant ecosystem around the world and are characterized by large spatial heterogeneity [1]. Urban landscapes exhibit rapid and pronounced environmental variation, such as flooding events from extreme rainfall and sensible heat rises with temperature increases of 2–10 °C [2,3]. Urbanization has also sharpened heterogeneity in population density and socio-economic conditions, exacerbating inequalities [4, 5]. Pronounced socio-economic inequalities are evident in the unprecedented scale of the vast informal settlements of low and middle-income cities [6,7]. Although such heterogeneity is expected to have important consequences for the spatio-temporal population dynamics of vector borne diseases, the joint effects of climatic and socio-economic conditions remain poorly quantified, especially for whole cities and across different spatial scales [8, 9].

Urbanization can alter both ecological and physiological parameters of mosquito vectors, altering the spread and emergence of mosquito-borne diseases. It can also improve infrastructure and environmental health, leading to better health care provision [10]. Studies of urban malaria have focused on Africa where the disease remains a predominantly rural problem, as the vectors are adapted to breed in rural environments [11, 12]. A common view is that urbanization reduces malaria transmission in African cities, given limited suitable breeding sites for the vectors [13], improved access to health care services and an increased ratio of humans to mosquitoes [12]. Nevertheless, transmission continues to persist in cities, and in some cases at even higher levels than those of surrounding rural areas [14].

In contrast to Africa, the Indian subcontinent harbors a truly urban vector, *Anopheles stephensi*, enabling the transmission of urban malaria for both parasites, *Plasmodium falciparum and Plasmodium vivax*. The mosquito breeds in various artificial containers within homes and construction sites [15, 16]. Given the rapid urbanization of this region, there is growing interest in the spatio-temporal structure of urban malaria and how it reflects the socio-economic and environmental heterogeneities of large cities [17, 18]. A previous study addressed malaria risk in the city of Ahmedabad in Northwest India [17]. The statistical models considered climate factors averaged across the whole city, and other drivers only at the mesoscale of districts. Thus, the role of local population density, humidity and high temperatures remain largely undescribed for whole cities and across spatial resolutions [16]. The recent reports of *A. stephensi* in the Horn of Africa raises concerns on the possible future expansion of urban malaria beyond its current geographical distribution [19].

Climate variability and climate change are expected to impact urban areas in particular ways, acting as major determinants of global health [20]. Some researchers have argued that at coarser spatial resolutions (e.g. >10 km), the effect of climate in urban areas may be negligible [21]. Others have argued that even at coarse aggregated scales, the effects of climate change should be evident, not only in cities, but also in large peri-urban areas [22]. The sensitivity of mosquito vectors to environmental variation at fine spatial scales has been studied [23, 24], with expected complex interactions with local features, such as housing density and material, vegetation cover, and distance to water [25, 26]. Since climate variables at coarse resolutions represent averages over large areas, land types and populations, they tend to hide extremes and can lead to spurious correlations with disease risk [27, 28].

Here, we use a space-time modelling approach to investigate the spatial distribution of urban malaria risk and the influence of both climatic and socio-economic drivers in the city of Surat, where an extensive surveillance program allows consideration of spatio-temporal variability at three different resolutions. We then discuss implications of these findings for malaria control and elimination efforts in the Indian subcontinent.

## Materials and Methods

### Study site and data description

The city of Surat presents ideal characteristics for our study, given the pronounced environmental and socioeconomic disparities, as well as a well-established surveillance program. The city is located on the banks of the Tapi River in the western part of India in the state of Gujarat (Fig 1). It is one of the fastest growing Indian cities due to immigration from various parts of Gujarat and other Indian states. Its unprecedented growth in the last four decades has led to a 10-fold increase in population [29]. It exhibits semi-arid weather with summer temperatures ranging from 37 to 44 °C with temperatures down to 22 °C in winter and averages of 28 °C during the monsoons. Rainfall ranges from 950 to 1200 mm per year (S1 Fig 1), and 90% of rainfall falls during the monsoon season, from June to September [30].

**Fig 1.**
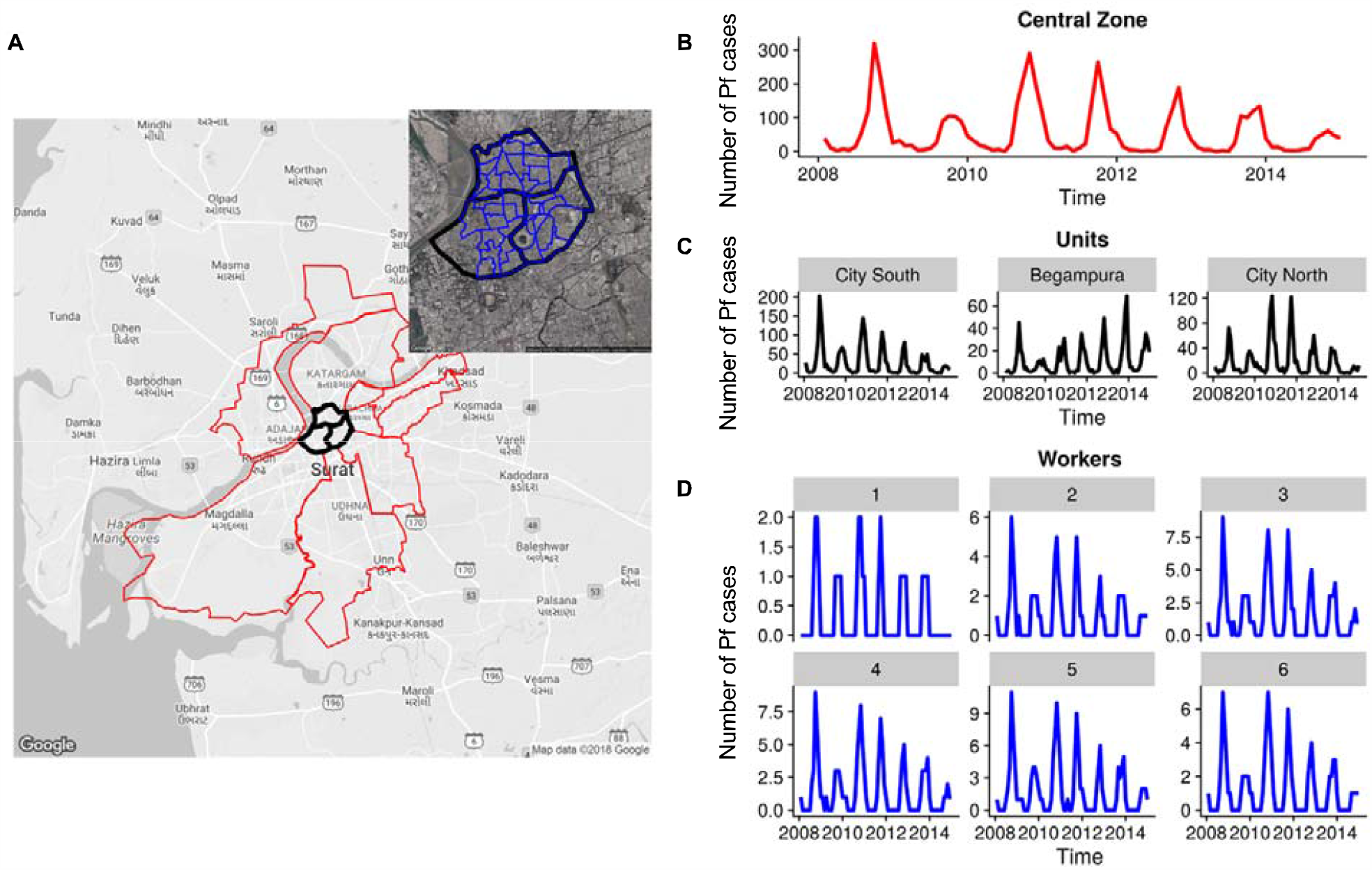
Location of the study area. (A). The 7 administrative zones of the city are depicted in red, and for illustration selected units (of a total of 32) are also shown in black within the central zone. Embedded within these, the subplot in the upper right illustrates the smallest reporting scale, that of the worker units. Time series of *P*.*falciparum* cases for (B) the central zone, (C) the three units and (D) the six worker units within the central zone.

In this largely semi-arid state where malaria is seasonally epidemic, peak incidence occurs between September and November [31, 32]. Surat exhibits an average of 1978 total cases per year (0.17 cases per 1000 population) for *Plasmodium falciparum*, and 6114 cases per year (0.36 cases per 1000 population) for Plasmodium *vivax* (S1 Fig 2). Malaria in northwest India, and in the state of Gujarat in particular, is seasonal and transmission is considered low and epidemic. The burden of malaria has been greatly reduced in the city since the epidemic emergence of the 1990s for both parasites. We focus our analyses on *P. falciparum* because this parasite remains the main target of control efforts. Dedicated control efforts (indoor residual spraying, breeding sites detection or insecticide-impregnated bed nets to prevent transmission) by the SMC (Surat Municipal Corporation) have kept malaria incidence at relatively low levels but have not eliminated the disease. Malaria still exhibits inter-annual variability with seasonal outbreaks of varying size (S1 Fig 2).

We obtained malaria data from 2004 to 2014 from the SMC disaggregated at three different levels of resolution corresponding to 7 zones, comprising 32 units, which are further divided among 478 ‘worker’ units. Epidemiological surveillance, active and passive, is conducted daily at the worker unit level (malaria control personnel is assigned to one unit of approximately 10,000 population). Active case detection (ACD) is performed through house to-house surveys by the malaria workers with each house visited every 30 days. The main responsibilities of the malaria workers are to identify fever cases in the last 15 days in a given household, to collect blood slides if a fever case occurred, and to provide radical treatment through follow-up visits. For passive case detection (PCD), each worker collects the number of infections reported by the relevant Urban Health Centers, with 32 such centers distributed homogenously across units in the city. The urban health centers collect and process the cases from the corresponding unit, with diagnoses conducted via blood smear microscopic examination.

We collated a multi-sourced spatio-temporal (climatic and socioeconomic) dataset. We reconciled data of different types and aggregation levels (economic, demographic) to the gridded climate data to generate three databases of malaria cases and covariates at the different levels of aggregation.

The socio-economic data is based on the 2010 census and a household survey at 90 locations and 400 households. Further data on population and slum density was obtained from the SMC. Climate data for temperature and Relative humidity was derived from satellite products. Land surface temperatures were extracted from both Terra and Aqua satellites of MODIS (https://earthdata.nasa.gov/eosdis/daacs/laads) for their overlapping period between 2008 and 2015. We also used these products to estimate surface relative humidity (RH) within the city. RH reflects the amount of moisture in the atmosphere. If RH is low, it will contribute to the drying of fuels; conversely, if RH is high, the fuels will absorb moisture from the atmosphere. To estimate surface RH in the region we used MODIS data, combined with some meteorological parameters obtained from ground-based measurements recorded at meteorological stations. MODIS data were processed to calculate precipitable water vapor (PW), and to extract surface air temperature in the clear sky. Finally, to reconciled different levels of aggregation an ordinary kriging method was used to generate an estimated interpolation surface for each covariate detailed description of the generated datasets is explained in Supplementary text 2.

## Data analyses

To examine spatial patterns of malaria risk independently from the inter-annual variation of incidence, we accumulated reported cases for a given year, and normalized this sum by the total yearly cases for the whole city. We then conducted univariate statistical analyses to evaluate the spatial dependency of the malaria cases (Fig 2 B and C). First, a univariate Moran Index (Moran’s I) was computed through time [33, 34] (Fig 2 C). Moran’s, I identify the global degree of spatial association (or how much the magnitude of an indicator in one location is influenced by the magnitude of the indicator in an area close to it) [34,35]. Then the Local Moran statistic was obtained to identify local clusters and spatial outliers [35] (Fig 2 A).

**Fig 2.**
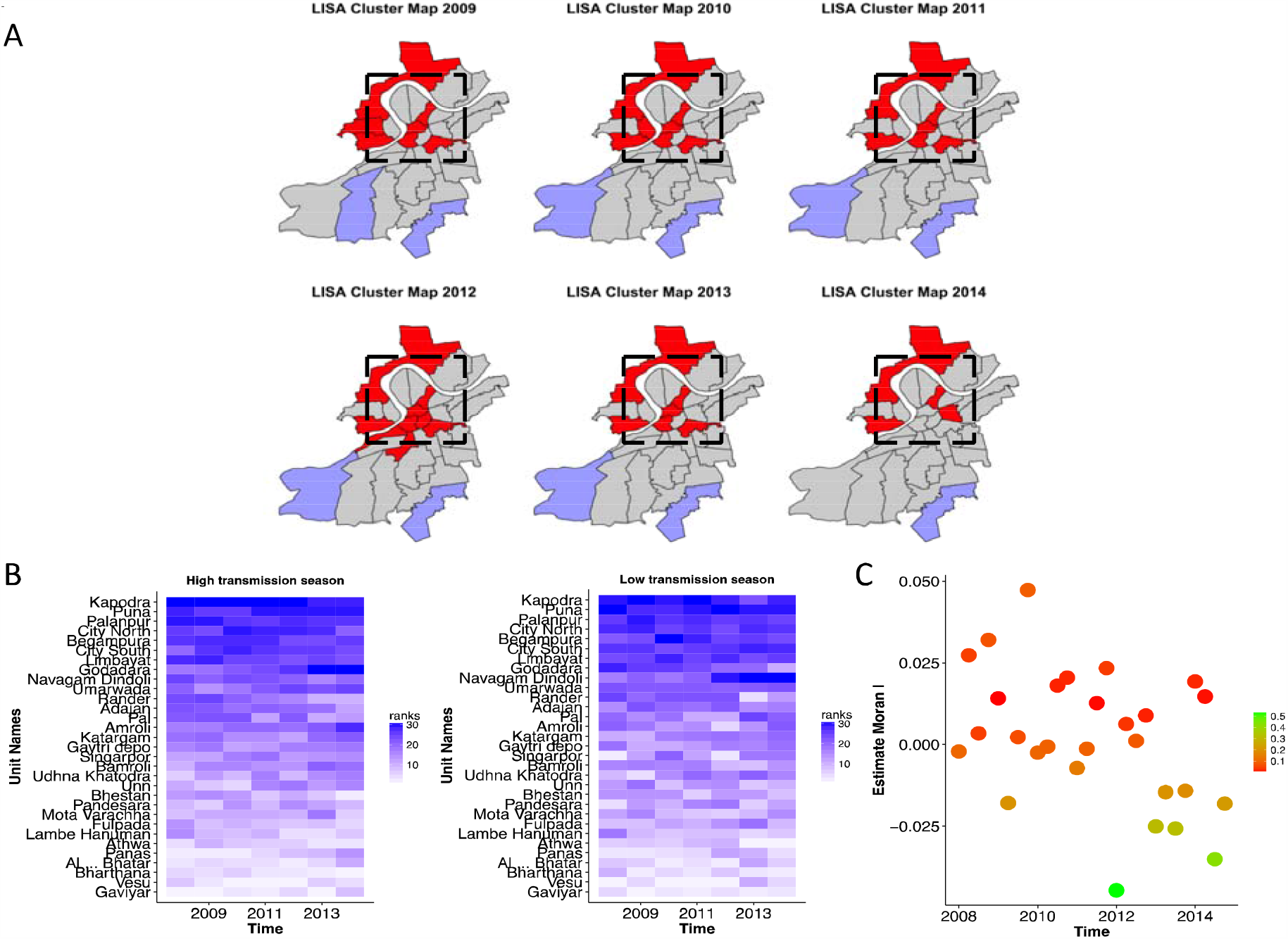
Spatial patterns. (A) Clusters in malaria incidence for the period 2008-14, identified using the local indicators of spatial associations (LISA) test/analysis. The resulting LISA cluster maps depict locations of significant Local Moran’s I statistics, classified by type of spatial association. This analysis specifically identifies units with spatial association in malaria incidence. Depending on the sign of the indicator (positive or negative), the local associations can be positive-positive, positive-negative, negative-positive or negative-negative. Positive-positive and negative-negative associations represent spatial clustering, where positive-negative and negative-positive correspond to spatial outliers and local spatial autocorrelation correspond to the core of a cluster (the actual “cluster” includes all the neighbors of a unit as well as the core. Here, red color depicts positive/positive associations (locations where high incidence is surrounded by high incidence) and light blue, negative/negative associations (locations where low incidence is surrounded by low incidence). Clusters are significant at p = 0.05 (based on 9999 permutations). The dotted box shows the units corresponding to the city center. (B) Distribution of cases normalized by population, with color intensity (from low [white] to high [blue]) corresponding to the ranking of incidence at the levels of the units. (C) The Moran’s I autocorrelation index is calculated bimonthly from 2008-2014. Color in the figures from red to green represents the significance of the spatial autocorrelation, red values are associations that are significant at 0.05 level.

To reduce the dimensionality of the socioeconomic variables (S1 Table 1) and to address the existence of a spatial pattern in these indicators, we used principal component analysis (PCA) to find the best low-dimensional representation of the variation in this multivariate data set (S1 Fig 3). Figure 2 A shows the organization of the variables in a low-dimensional socioeconomic space as well as in physical space at the level of units. We specifically considered the spatial distribution of the components accounting together for more than 80% of the variance in socioeconomic space. Figure 2b shows the spatial distribution of the PCA scores, corresponding to the new coordinates of a given unit in the space defined by the principal components. The maps of the scores for the three main components summarize how the variance in socioeconomic conditions is organized in space (Fig. 3 C-D). Then we evaluated the association between the values of these scores for each PC and the mean malaria cases across the different units (S1 Fig 4 B).

**Fig 3.**
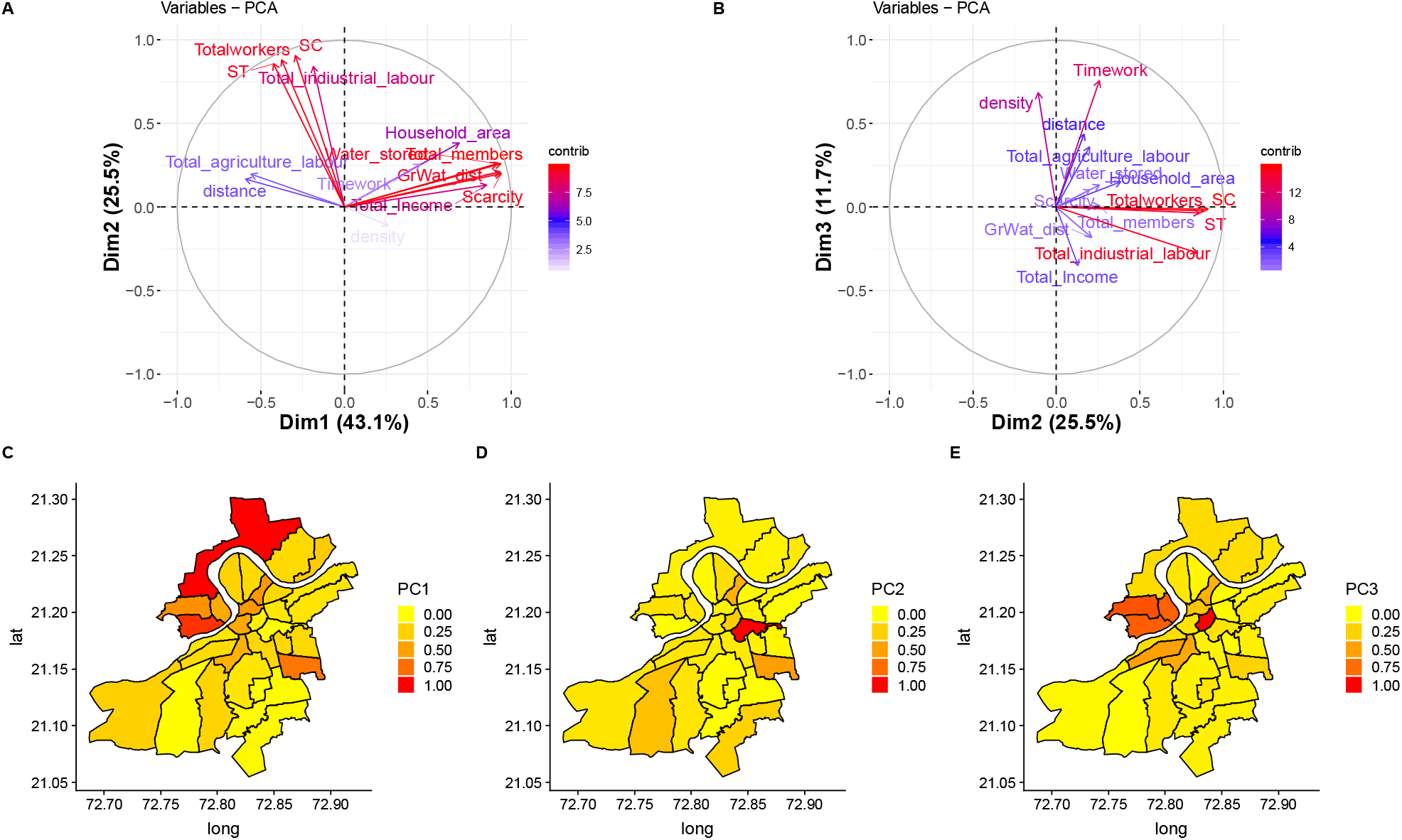
Principal Component Analysis: contribution of socioeconomic variables to the three main axes of variation, and spatial distribution of socioeconomic variation in these dimensions. Vectors in the top panel depict contributions of the socioeconomic variables to two of the principal components (PC) at a time, in A) the first and second, and (B) the second and third dimension. The colors from light blue to red show the respective contributions from low to high. PC1 largely represents economic level, PC2 is associated with labor and employment, likely representing the effect of movement and exposure in particular environments within the city, and PC3 exhibits a strong contribution from population density. The maps in panels (C), (D) and (E) show the distribution over the different spatial units of component scores for PC1, PC2 and PC3 respectively. These component scores correspond the new coordinates of a given unit for a given PC, with intensity from low to high in colors from yellow to red. We also evaluated the association between the values of these scores for each PC and the mean malaria cases across the different units (S1 Fig 4 B),

### Statistical models

A hierarchical model framework was applied to assess the relative contribution of climatic and socioeconomic factors in determining space-time urban malaria case patterns. Generalized linear mixed models (GLMM) were formulated including climate covariates, PCA terms, and random effects (to account for spatial dependency structures, seasonality, and interannual variability attributed to unobserved factors, such as control efforts, quality of health care services, and local health interventions). The general form of the GLMM is as follows:

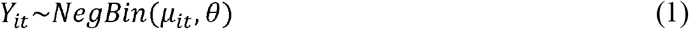

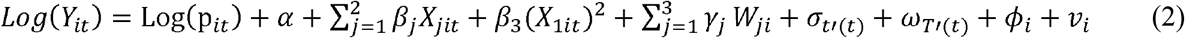

where *Y*_*it*_ are the malaria counts for each administrative unit (where i = 1,…,32 for the unit level, i = 1,….,7 for the zone level and i = 1,…,486 for the worker unit level). The population per 1000 p_it_ is treated as an offset. The variablesh*X*_*jit*_ represent the selected climate influences: temperature one month earlier (j=1), and humidity two months earlier (j=2). These significant lags where identified using the Autocorrelation Function (ACF) and the Partial Autocorrelation Function (PACF) (S1 Fig 5). Quadratic terms in these variables were included to allow for nonlinear effects, in particular, documented empirical declines in response to temperature after a given maximum. To incorporate socioeconomic factors in the model, we used their principal components [36]. This allows consideration of fewer factors that effectively summarize the main variability in the survey data, and also avoids the problem of correlations between the original socio-economic variables. The variables *W*_*ji*_ correspond respectively to the components PC1 (j=1), PC2 (j=2) and PC3 (j=3) identified by the PCA analysis. To account for the effect of seasonal variation we added a monthly random effect σ_*t*′ (*t*)_ *t*′ (*t*) = 1, … .12 (this monthly random effect has a random walk prior to allow for the dependency of a given month on the previous month). Moreover, to account for interannual variation, a yearly random effect was introduced, with ω_*T*′(*t*)_ *where T* ^′^ (*t*) = 1 … .11. Finally, spatially unstructured random effects were included via *ϕ*_*i*_, to account for over dispersion and unknown and unmeasured effects. Spatially structured effects were included to allow for spatial dependency between neighboring areas, via *v*_*i*_ (with a conditional autoregressive CAR, prior) [37, 38] (For details, Supplementary text 3).

### Scale dependency

Based on the best model identified at the intermediate level of aggregation of units, we then tested how the magnitude and significance of the different factors changed at coarser or finer resolutions. We used the best model structure identified for the 32 units and re-fitted the model at a higher (478 worker units) and lower resolution (7 zones), following the same methodology described in Supplementary text 3. We also evaluated changes in the 95% credible interval (CI) of the different estimated parameters to determine whether it contained zero (Table 3).

### Model assessment

To assess the predictive ability of the best fitting model at the intermediate spatial scale (hereafter, the intermediate model), posterior predictive distributions of malaria incidence were obtained for each unit and month. To summarize this information, the observed and posterior predictive mean malaria risk estimates were aggregated across space, and predictions for each unit in high and low incidence years were generated (Fig 4 A-B). We compared the temporal evolution of the fitted posterior median cases with the observed cases for Surat as a whole (S1 Fig 6).

**Fig 4.**
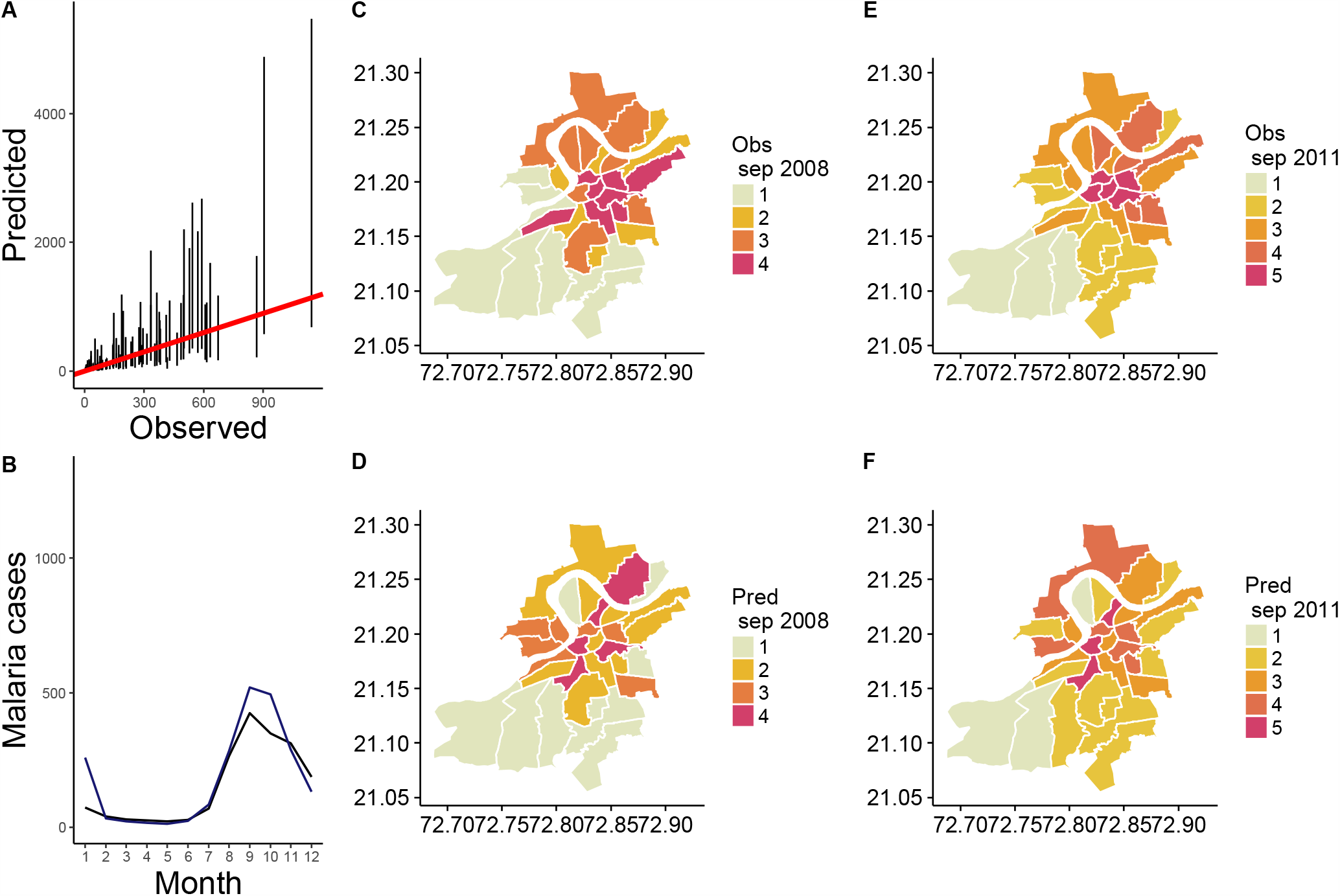
Observed versus predicted *Plasmodium falciparum* (PF) cases. A) Total malaria cases. The identity line (in red) is shown for comparison. B) Seasonal pattern for the observed cases averaged over all years (red) and the median of 1000 model simulations (blue). The 2.5% and 97.5% percentiles of the simulated data are shaded in light red. Comparison of observed and predicted cases for 2008 (C-D) and 2011 (E-F). The colors in the maps progress from yellow to red based on quantiles generated by considering all zero cases in one class, and by subdividing all remaining non-zero cases into four intervals. The resulting 5 categories correspond to no cases (1), very low (2), low (3), high (4) and very high (5) cases. Comparisons of the quantiles in the maps reveal that 73% and 66% of their values correctly match for years 2008 and 2011, respectively.

We also compared the proportion of places in which the intermediate model accurately predicts malaria incidence. For this, we classified the cases into 5 categories, generated by considering all zero cases in one class and by subdividing all remaining non-zero cases into four equally sized intervals. The resulting categories correspond to no cases, very low, low, high and very high cases (based on different quantiles of the cases distribution). We then mapped the categories of the spatial predictions and observed cases for 2008 (Fig 4 C-D) and 2011 (Fig 4 E-F), quantifying the number of times the values correctly matched.

## Results

The pattern that results from ranking malaria risk based on incidence is largely stationary, as shown in Figure 1 and S1 Fig 7, with locations of high and low risk persisting over time, independent of the inter-annual variation in total malaria cases. This regular spatial pattern suggests the existence of strong underlying spatial determinants, that are themselves largely stationary over the temporal scales of malaria variation considered here (Fig 2 A and S1 Fig 7). The stationary pattern in malaria is supported by a significant spatial autocorrelation through time (Fig 2 C). Results from the local indicators of the spatial association test (LISA) show that the units spatially associated with high malaria risk differ significantly between the center of the city and its periphery (Fig 2 B). Specifically, units in the central and northern part of the city exhibit (positive/positive) associations in malaria risk or hotspots, whereas some units in the periphery show (negative/negative) associations or coldspots. The clusters identified through spatial associations also show a striking regularity from season to season (Fig 1 B).

For the socioeconomic data from census and surveys in the city (S1 table 1), we first assessed if we could simplify the variation by summarizing it in a low dimensional space. The results from PCA showed that the first three dimensions can explain 80.9% of the variability among units. Dimensions 1, 2 and 3 respectively explained 45.1%, 27% and 3, 8.84% of the variance (Fig 3 A-B). These three dimensions were therefore retained for further analysis, to include socioeconomic covariates in the spatio-temporal statistical model via a low number of covariates. Figure 3 shows the loadings or contributions of each socioeconomic variable to the different principal components, plotted in each of the two-dimensional subspaces PC1-PC2 in (A), and PC2-PC3 in (B). These contributions are also given in S1 table 2. They show that PC1 largely represents economic level and is also correlated with the amount of water stored. PC2 is associated with labor and employment, likely representing the effect of movement and exposure in particular environments within the city, and PC3 exhibits a strong contribution from population density. Panels C-D of figure 3 show the spatial distribution of the PCA scores for the different units for each of the three principal components. These scores are the coordinates of the units in each of these three dimensions. They show that the spatial pattern for economic level (PC1) and population density (PC3) closely matches that of disease risk (compare Fig 3 C-D and S1 Fig 7).

We then explored if malaria risk is associated with the space-time variation in environmental factors (relative humidity RH, and temperature) and with the spatial variation in socioeconomic variables, including population density, summarized in the three main components of the PCA (Fig 3). S1 Fig 4 A illustrates the temporal associations between ranked malaria risk and mean temperature and humidity, respectively. A statistically significant linear correlation was found only between malaria risk and relative humidity (p < 0.01). S1 Fig 4 B shows significant spatial correlations (p<0.05) between the principal components 1 and 3 and the mean ranked cases. These exploratory analyses provide an indication on the covariates that might contribute significantly in a spatio-temporal statistical model of the variability in cases across the city. To more formally select those variables, we proceeded to consider different GLMM models of increasing complexity.

Goodness-of-fit metrics for the GLMM models are shown in Table 1. Models that include or neglect, the effect of given climatic and, economic/demographic variables, as well as neighborhood structure, are compared on the basis of WAIC (Watanabe-Akaike information criterion). The selected model that best accounts for the spatio-temporal variation in malaria incidence included the combined effects of temperature, humidity, principal components 1 and 3, and seasonal and spatial random effects. Specifically, accounting for these factors in combination to the random structured and unstructured effects, explains 58% of the variance based on the R^2^ likelihood ratio test (Table 1). Of the 58% total variance explained, 41% is explained by the unstructured spatial random effect, monthly (e.g. seasonality) and yearly random effects and the Structured random effect. An additional 7% is accounted for by principal components 1 and 3, and an additional 11%, by climate factors (Temperature and Relative humidity). To examine whether the selected climate variables better explain malaria risk at the regional or local level, we compared models with temperature and relative humidity that were either spatially resolved or averaged over all the spatial units at the global level of the city. The best fitting model incorporated aggregated humidity and spatially-explicit temperature. These findings suggest that temporal variation in malaria cases across years is explained by the interannual variability in humidity, whereas the spatial variation in cases is influenced local temperature heterogeneity within the city.

**Table 1.** Comparison of goodness of fit. Comparison of goodness of fit for the different models tested based on different model selection criteria (DIC, WAIC and R^2^ based on a likelihood ratio test). Global humidity or temperature correspond to averages over all the units; whereas spatial humidity and temperature indicate their local value for the 32 units (smaller DIC and WAIC values indicate better fit).

| *Model* | **DIC** | **$R^2_{LR}$** | **WAIC** |
| --- | --- | --- | --- |
| *Baseline: month (e.g. seasonality) and year random effects* |  |  |  |
| (A) Unstructured spatial random effect, monthly (e.g. seasonality) and yearly random effects +<br>*Structured random effect* | 4960 | 0.41 | 810.7 |
| *Baseline + socio-economic (different combinations of PCA1, 2 and 3)* |  |  |  |
| *(B) Baseline + PC1* | 4947 | 0.43 | 803.7 |
| *(C) Baseline + PC1 + PC2* | 4949 | 0.42 | 806.4 |
| *(D) Baseline + PC1 + PC2 + PC3* | 4942 | 0.44 | 803 |
| *(E) Baseline + PC2 + PC3* | 4943 | 0.44 | 803.1 |
| *(F) Baseline + PC1 + PC3* | 4936 | 0.48 | 801.5 |
| *Baseline + socio-economic + different combinations of climate factors* |  |  |  |
| *(G) Baseline + socio-economic + spatial temperature* | 4931 | 0.5 | 798.2 |
| *(H) Baseline + socio-economic + averaged temperature + averaged humidity* | 4928 | 0.53 | 796 |
| *(I) Baseline + socio-economic + spatial temperature + averaged humidity* | 4922 | 0.58 | 792.6 |
| *(J) Baseline + socio-economic + averaged temperature + spatial humidity* | 4930 | 0.51 | 799.7 |
| *(K) Baseline + socio-economic + spatial temperature + spatial humidity* | 4927 | 0.53 | 796.3 |
| *(L) Baseline + socio-economic + spatial polynomial temperature + spatial humidity* | 4928 | 0.52 | 795 |

S1 Table 3 summarizes the posterior mean parameter estimates for the best model. All parameters are significantly different from zero, with posterior distributions from the two chains well mixed and converged based on the Gelman-Rubin diagnostic (S1Table 3). PCA 3 exhibits a positive and statistically significant association with malaria incidence. The effect of temperature is negative, consistent with the non-monotonic effect of temperature on the reproductive number R_0_ for malaria, where the relationship changes from positive to negative as temperatures increase beyond the thermal maximum toward the high end of the temperature spectrum [39,40]. Here, temperatures lie above this thermal maximum, the reason why only a negative effect is identified whereas consideration of a polynomial term did not improve the fit. This negative effect is consistent with increases in temperature above a certain threshold influencing mosquito and parasite physiological and demographic parameters in a way that decreases transmission intensity [40]. Finally, relative humidity shows a significant and important enhancing effect on malaria risk across the city.

A comparison of predicted and observed malaria cases is presented in Figure 4. There is in general a good agreement in time, with a tendency to underpredict the largest peaks. The maps show spatial comparisons for observed and predicted cases at the unit level for representative years, namely 2008 (for high incidence years) (Fig 4 C and D), and 2011 (for low incidence years) (Fig 4 C-F). In general, the predicted patterns reflect the observations for individual units (Fig 4 C-F) and for averages over the units (Fig 4 A-B). We observe correct quantile predictions for 73% and 66% of the units respectively (S1 Table 4). Overall our GLMM model tends to under predict malaria, especially for the city center in years with low epidemic peaks such as 2008. However, for years with large epidemics like 2011, the model is able to capture the spatial pattern, including across the southeast and northwest of the city.

To compare the coefficients of the model fitted at different spatial levels (zones, units, workers), we refitted the intermediate (unit) model for 2008 to 2014 (the period of time for which we also have high resolution malaria data) at both the aggregated zone level (7 zones) and the disaggregated workers’ level (478 locations). Table 2 shows changes in the values of the model coefficients and their level of significance (see Table 2) at the different spatial scales. (Parameter estimates are considered to be statistically significant if their 95% credible interval does not contain zero). Temperature and humidity show variation in their contribution across the spatial levels: when we disaggregate the system, the effect of temperature strengthens whereas the effect of humidity weakens. Also, the effects intensify for PC1 and PC3 (largely reflecting respectively level of income/economically deprived communities, and population density), both for the value and the significance of the corresponding coefficients from lowest to highest resolution.

**Table 2.** Model comparisons. Comparison between the parameter estimates from models fitted at the three levels of aggregation (zone, unit and workers unit). If the CI did not contain zero, the corresponding covariate contributes significantly to the model fit. Bold numbers indicate statistically significant parameter estimates (i.e. the 95% credible interval does not include zero).

| *Level of aggregation* | *Temperature* | *Relative humidity* | *Principal component 3* | *Principal component 1* | *Monthly random effect* | *Yearly random effect* | *Spatial unstructured parameter* | *Spatial structured parameter* |
| --- | --- | --- | --- | --- | --- | --- | --- | --- |
| *Zone (n = 7)* | [-0.054,0.132] | **[0.614, 1.028]** | [0.112, -0.098] | [-0.115, 0.02] | **[0.621, 0.742]** | **[0.694, 1.101]** | **[0.722, 1.625]** | [-0.123, -0.298] |
| *Ward Unit (n = 32)* | **[-0.139, -0.307]** | **[0.506, 0.946]** | **[0.039, 0.259]** | **[0.369, 0.723]** | **[0.234, 0.556]** | **[0.9674, 2.025]** | **[0.681, 1.416]** | **[1.681, 2.416]** |
| *Worker unit (n = 478)* | **[-0.214, -0.416]** | [-0.06, 0.892] | **[0.134, 0.398]** | **[0.401, 0.876]** | **[0.682, 0.803]** | **[0.422, 0.532]** | **[0.543, 0.923]** | **[1.724, 2.191]** |

## Discussion

Urban environments exhibit pronounced heterogeneity from rapid and unplanned urbanization [18, 41, 42]. Our results underscore the importance of considering this spatial heterogeneity for predicting urban malaria risk in the Indian subcontinent. The models presented here build upon the results of Santos et al. (2016) for a different, inland, city of Northwest India, by addressing the joint role of socioeconomic, demographic and climatic factors in determining malaria risk at different spatial scales. We find that the two climate covariates contribute differentially to explain malaria variation: humidity helps in capturing inter-annual variability and peak timing of outbreaks; temperature explains some of the spatial variation synergistically with socio-economic covariates. These effects are confirmed when we fit the intermediate model structure at the two other resolutions, whereby the variation explained by temperature is highest for the finer granularity and that explained by humidity increases at the coarser one.

These comparisons were performed on the basis of the model structure selected at the intermediate spatial level, and not via a systematic model selection analysis at the three spatial scales. We note however that the independent variables selected at the intermediate level include many of the initial factors interrogated. In particular, the intermediate model selected includes socio-economic variables and population density (through PC1 and PC3), climate covariates (temperature and humidity), and structured and unstructured random effects. Thus, when examining the significance and values of the model coefficients at the higher and lower resolution, we are effectively addressing how the corresponding variables contribute to the spatio-variability of cases across resolutions. It is of course possible but highly unlikely that a completely different “best” model would have been selected, had we performed model selection at each of the levels, rather than identified the significant coefficients remaining. Thus, the comparisons of the models fitted at the different levels likely reflect the importance of socio-economic and climate factors.

Our Principal Component Analysis shows the presence of three distinct explanatory components summarizing most of the variability across spatial units in the socio-economic covariates. The first component largely summarizes the important effect of economic disparities which can modulate host exposure to mosquitoes, and of water management, which would influence the recruitment of these vectors [43]. Specifically, for India access to water is an important determinant of malaria risk, given that water is supplied irregularly, leading to water storage within houses, which in turn creates multiple breeding sites for the mosquito in overhead tanks, cisterns and cement tanks [44]. The second component largely corresponds to variation in labor and employment, which likely relates to human mobility [45]. Movement of the vectors themselves is expected to act at very local scales since experiments have demonstrated that mosquitoes do not travel very far [46] and typically stay within the same residence for days [40]. Finally, the third component represents largely variation in population density which is critical to epidemic spread, particularly in urban landscapes with pronounced heterogeneity in population distributions. Population density can influence the individual risk of infection though its effect on the local abundance of mosquitoes [45].

Here, population density (represented by PCA3) was shown to positively influence malaria risk. This finding opposes the typical expectation from mathematical models for the transmission of the disease in which the rate of individual infection (the force of infection) decreases with human population size [45]. For urban malaria, higher population density could result in higher water storage concentrations in close proximity to people. Our finding is consistent with that of Romeo-Aznar et al. (2018) who recently proposed that vector abundance for dengue can increase with human population density through higher mosquito recruitment, especially in poor areas of cities. The authors found that in poor areas of Delhi an increase faster than linear is sufficiently fast to generate a positive trend in the force of infection with higher human population.

Our study further shows a negative relationship between malaria cases and high temperatures. This complements the more commonly documented positive relationship near or below optimal temperature conditions for physiological and epidemiological parameters of the mosquito, and for the malaria parasite within the mosquito [39]. Temperatures in Surat are generally higher than those identified as optima for malaria vectors. The negative relationship at high temperatures, beyond this optimum, emphasizes the need to better understand the high end of the temperature spectrum for vector-borne infections, the least studied part of physiological curve [38, 39]. Moreover, humidity and temperature show contrasting effects in our model in terms of whether they act at local or global city scales. A model that incorporates aggregated humidity (averaged for the whole city) and spatially-explicit local temperature, performs better than one with both climate variables spatially resolved. This difference could be explained by the strong dependence of humidity on winds, which can alter evaporation by changing water vapor in the air. Because winds tend to vary at a regional scale, humidity would also manifest change over large distances [47]. Conversely, temperature can exhibit large variation within a city at the local level, given the pronounced heterogeneity of impervious surfaces, with different radiative, thermal, aerodynamic, and moisture properties [48, 49]. This conclusion should be further examined with a higher number of local humidity measurements on the ground to cross-validate satellite-derived values. Similarly, a higher number of households could be included to describe socio-economic variation across the city.

Despite these limitations, our spatio-temporal statistical model captures the seasonal pattern and the main trends in the inter-annual variation in malaria cases. Our model is also able to predict the spatial variation in epidemic peaks. The model framework proposed here could be further improved by including: mobility fluxes derived with movement models from the spatial distribution of the population, replacing the near-neighbor effects on transition probabilities; other environmental heterogeneities such as river discharge and soil moisture. Temporal changes of the city structure itself would also be informative, including changes in the local speed of urbanization and the development of informal settlements. A better understanding of the effect of population density on vector abundance is another important consideration.

Surat has experienced strong malaria interventions in the last three decades reflected in a negative trend in the number of reported cases from the 1980s and 1990s to the 2000s. Since 2002 the city has experienced seasonal outbreaks with the overall level of malaria incidence remaining fairly constant. The stationary pattern of spatial risk described here, together with major drivers of this variation, indicate that targeted control could help reduce transmission even further, and that control measures could be implemented ahead of the season based on known spatial heterogeneity. In particular, ongoing efforts to provide better access to water may contribute to reduce transmission of urban malaria, and possibly that of other vector-borne infections. Although we were unable to separate here the correlated effects of poverty and water access/storage, this is an important area for further study. Ultimately, at longer time scales, a reduction in poverty concomitant with better access to an uninterrupted water supply is fundamental to reduce and eliminate malaria not only within cities but at a regional and national level. Also, a deeper understanding of humidity and temperature effects on malaria transmission would be key to predict future impacts of climate variability and climate change on the disease. Humidity and temperature are expected to increase under future climate projections for the Indian subcontinent [50], specifically in the northwest part of India [51, 52]. A better understanding of how climate factors influence malaria transmission within urban environments can inform India’s target of malaria elimination by 2030.

## Contributors

MP and MSV conceived and designed the study. MSV implemented the statistical analysis and wrote the first draft. MSV, MP, RL, LA analyzed the data and the results from the models. VK, VD and AK provided expertise on the data and on the epidemiology of the region. All authors read, contributed with literature search, edited and approved the final manuscript.

## Data sharing

The authors are open to sharing statistical codes and study data. Agreement of the Surat Municipal Corporation the data provider, will be required for any data sharing.

## Acknowledgements

We are grateful to the commissioner of Surat Municipal Corporation for providing the malaria data for the city. We also thank Dr. Menno Bouma for insightful discussions in early stages of this project, and Dr. Vimal Mishra from the Gandhinagar Institute of Technology for technical advice in the calculation and interpolation of the humidity data as well as the analysis of the climatological data and patterns. MP was supported by NIH (grant number 1R01AI153444). RL was supported by a Royal Society Dorothy Hodgkin Fellowship. We acknowledge the Gardner High Performance Computing (HPC) cluster from the Center for Research Informatics of the University of Chicago for computational resources.

