## Supplementary information for "Quantifying climatic and socio-economic influences on urban malaria in Surat, India: a modelling study"

**Supplementary Materials**

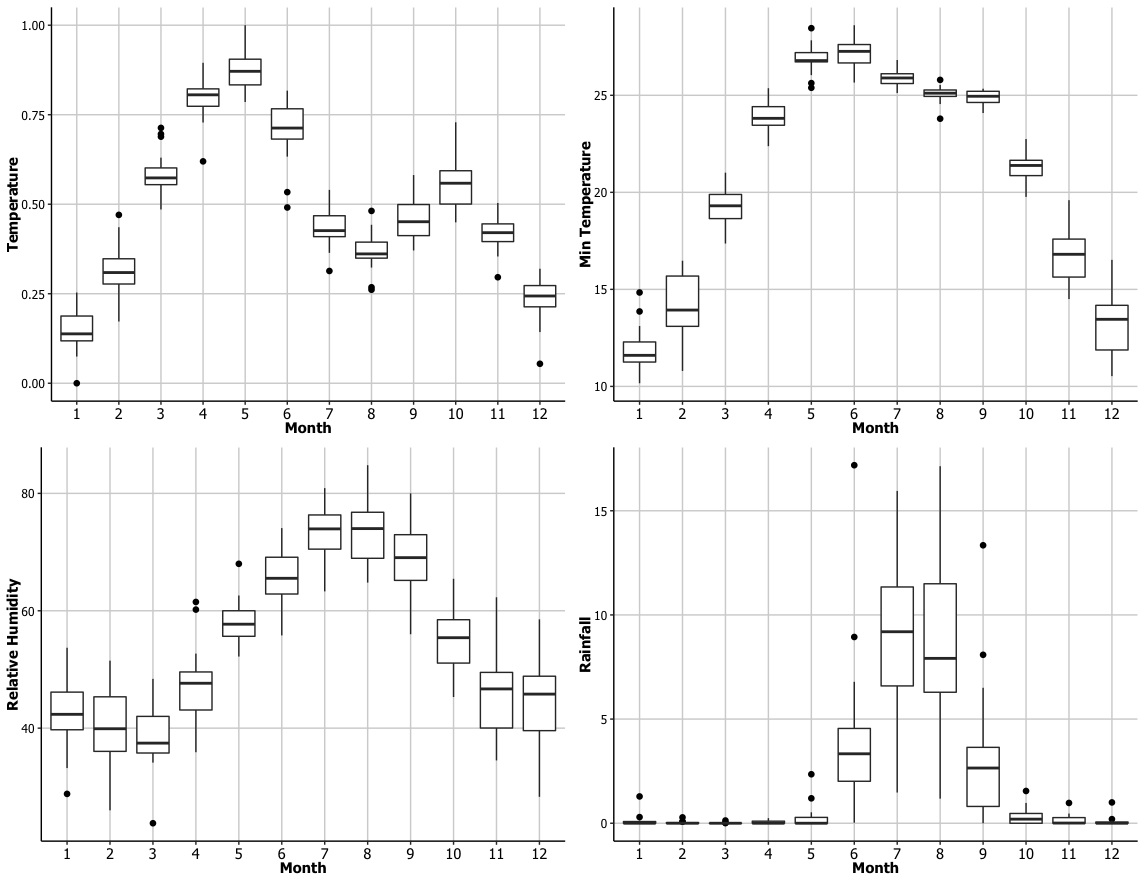

**Figure S1.1.** Monthly summaries of (a) mean temperature, (b) minimum temperature, (c) relative humidity and (d) rainfall in the city of Surat from 2008 to 2015, based on Surat station data from the India meteorological department (http://dsp.imdpune.gov.in/)

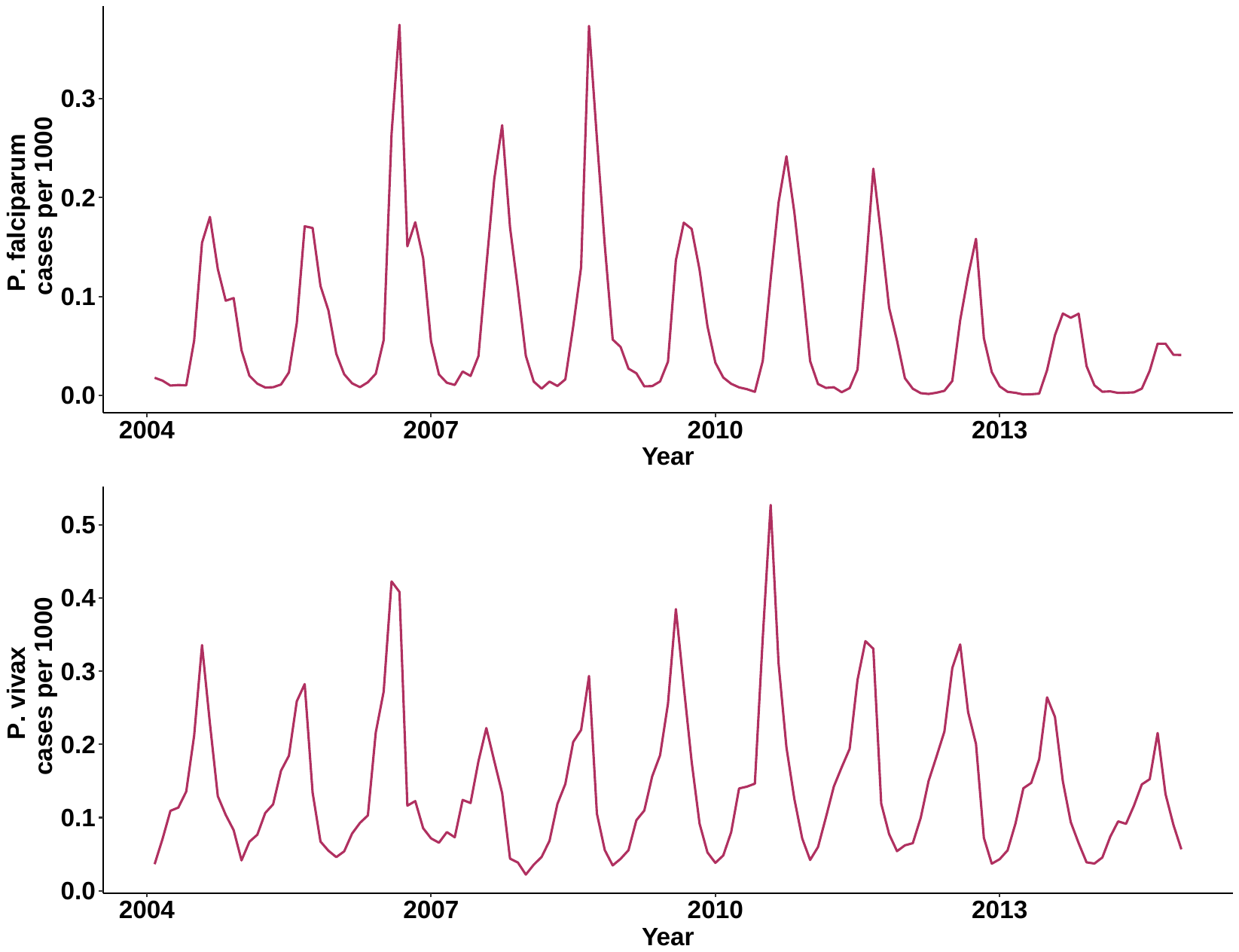

**Figure S1.2.** Monthly reported a) falciparum and b) vivax incidence per 1000 people in the city of Surat.

**Table S1.1.** Socioeconomic covariates included in the analysis

| **Variable** | **unit** | **Description** | **Justification** |
| --- | --- | --- | --- |
| Total industrial labor | persons | Number of people dedicated to industrial jobs | This variable is a proxy of employment in the city |
| Total workers | persons | The workers comprise 312 million main workers and 88 million marginal workers (defined as those who did not work for at least 183 days in the preceding 12 months to the census taking) | This variable is a proxy of employment in the city |
| Density | persons/area | This variable accounts for the number of human hosts per square kilometer in the city. | This variable is included to account for unobserved effects in the force of infection, including in particular increasing vector recruitment with human density as in Romeo et al. 2018. |
| Total Income | Rupees/per person | Per capita income in India is the mean income of the people in an economic unit. It is calculated by measuring all sources of income in the aggregate and dividing it by the total population. | We are including this variable since we expect this variable to correlate with disease risk and exposure. |
| Household area | Sq meters | This is the average house for each unit (ward). | We are including this variable since it’s a proxy of economic level in LMIC |
| Water stored | Cubic meters | This variable is considered because the vector breeds in water containers | Given that the vector breed within houses the amount of water stored is an important covariate. |
| Total members | persons |  |  |
| Distance to river | km | Proximity to de rivers |  |
| Total agriculture labor | persons | Amount of people dedicated to agricultural activities. | This variable could be a proxy for agricultural activities which might increase the availability of the pools of water with breeding sites for mosquitoes |
| SC (Scheduled Castes) | persons | officially designated groups of historically disadvantaged people in India. | This is a proxy of vulnerable communities (usually economically deprived) in the city |
| ST (Scheduled tribes) | persons | Indigenous people officially regarded as socially disadvantaged. | This is a proxy of vulnerable communities (usually economically deprived) in the city |
| Time to work | Hr/day | This is the amount of time that people commute | This variable is a proxy for the average commuting time for the people of each unit. |
| Distance to water sources | meters | This is the distance to main water bodies in the city | This distance can be a measure of exposure to breeding sites. |
| Scarcity | -- | This index account for the lack of water in some of the units | This variable provides an indirect measure for the amount of water that every unit store in its houses. |

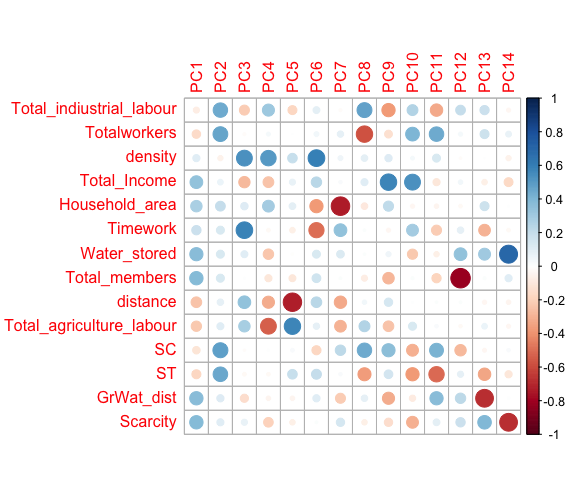

**Figure S1.3. Principal Component Analysis** scores (the transformed variable values in the new space defined by the principal components) or correlation coefficients between the variables (rows) and the principal components (columns).

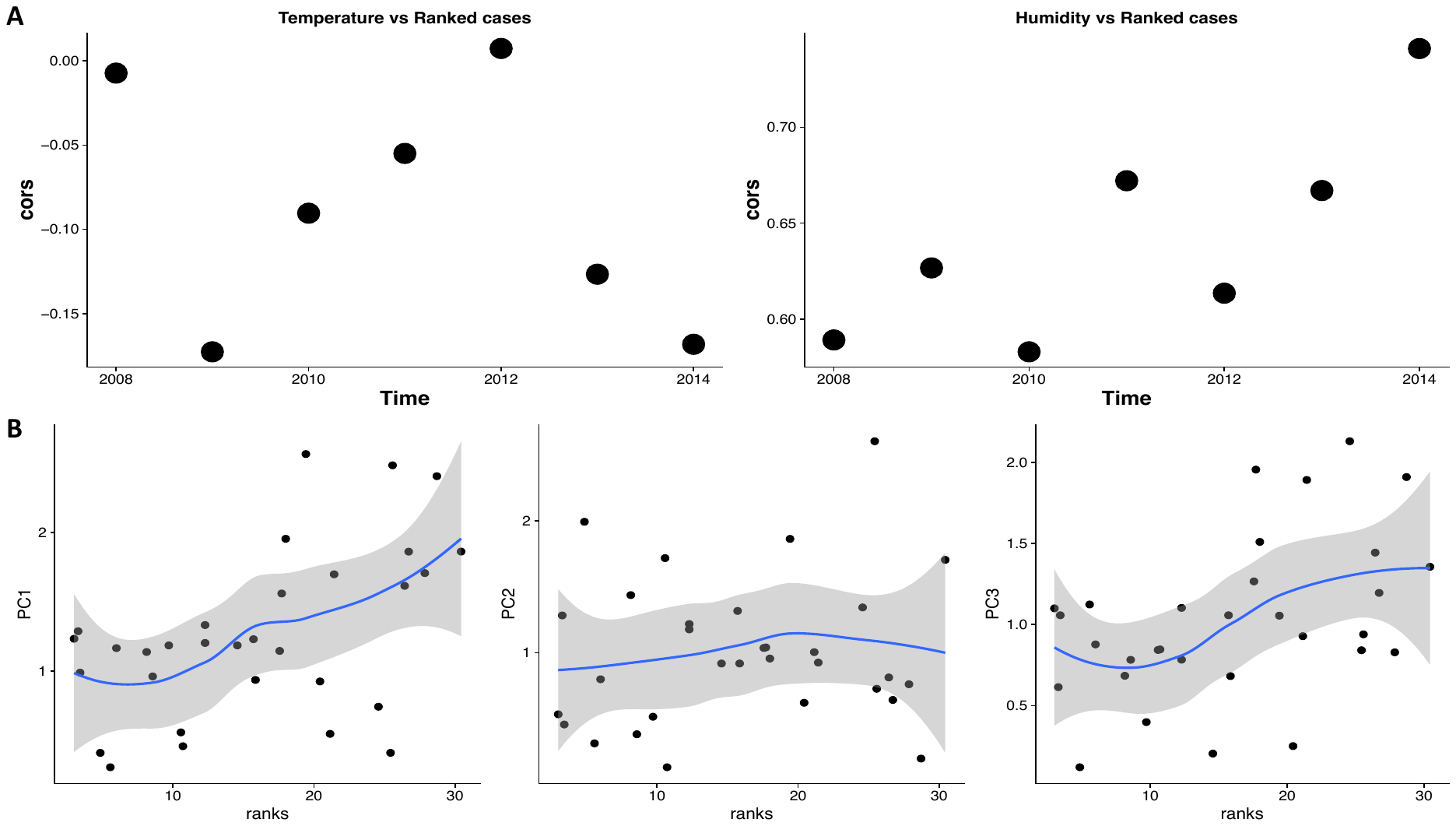

**Figure S1.4. Correlations between cases and drivers.** A) Top panels show temporal correlations plots between cases and temperature (R^2^ = 0.21, p > 0.01) and humidity (R^2^ = 0.72, p < 0.01), respectively. B) Bottom panels show correlation plots between the cases and the PCA scores of the 3 dimensions identified with the PCA in figure 2. These specifically show significant associations between cases and PC1 (R^2^=0.65, p < 0.05) and cases and PC3 (R^2^=0.53,p < 0.05).

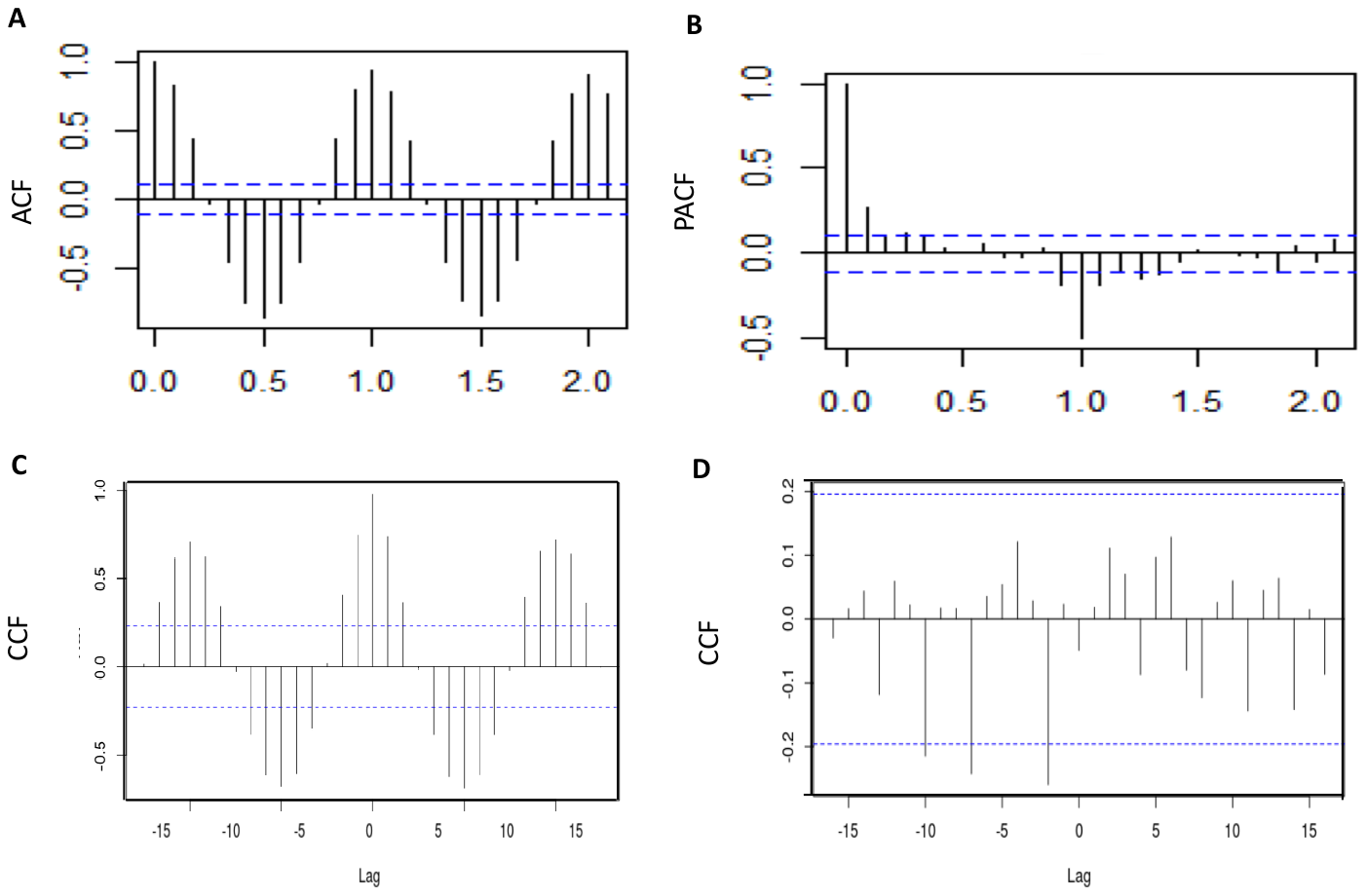
**Figure S1. 5. Autocorrelation Function (ACF), Partial Correlation Function (PCF) and Cross-Correlation Function CCF.** In (A) and (B) Show the ACF and PACF of monthly malaria cases respectively showing the presence of autoregressive process of order 1. In (C**)**. CCF between malaria cases and temperature with a significant lag of 2 months, and in(D), CCF between cases and humidity showing a significant lag of one month. Dashed lines represent the 95% confidence interval for correlations expected to arise randomly.

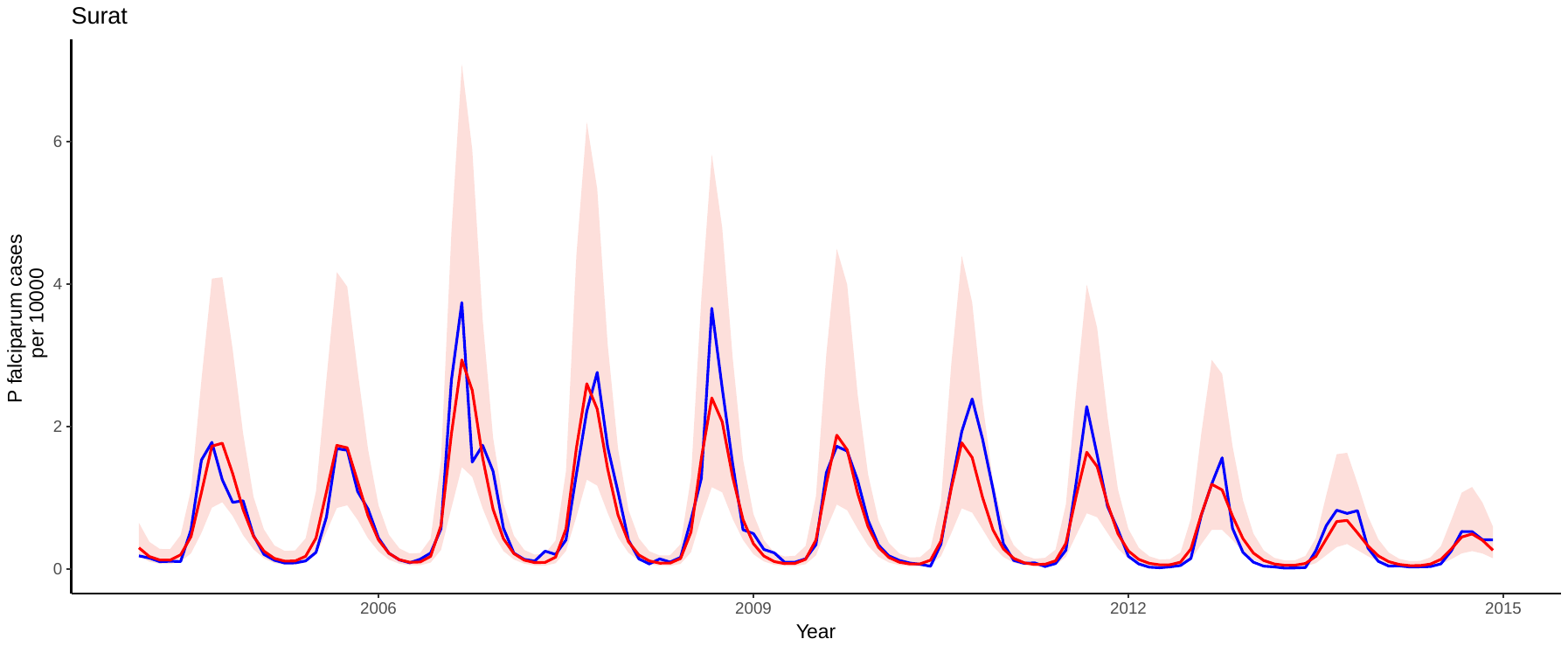

**Figure S1. 6.** Posterior median incidence (red curve) and 95% credible intervals (red shaded region) aggregated across all units from 2005 to 2014. Observed incidence is shown in blue.

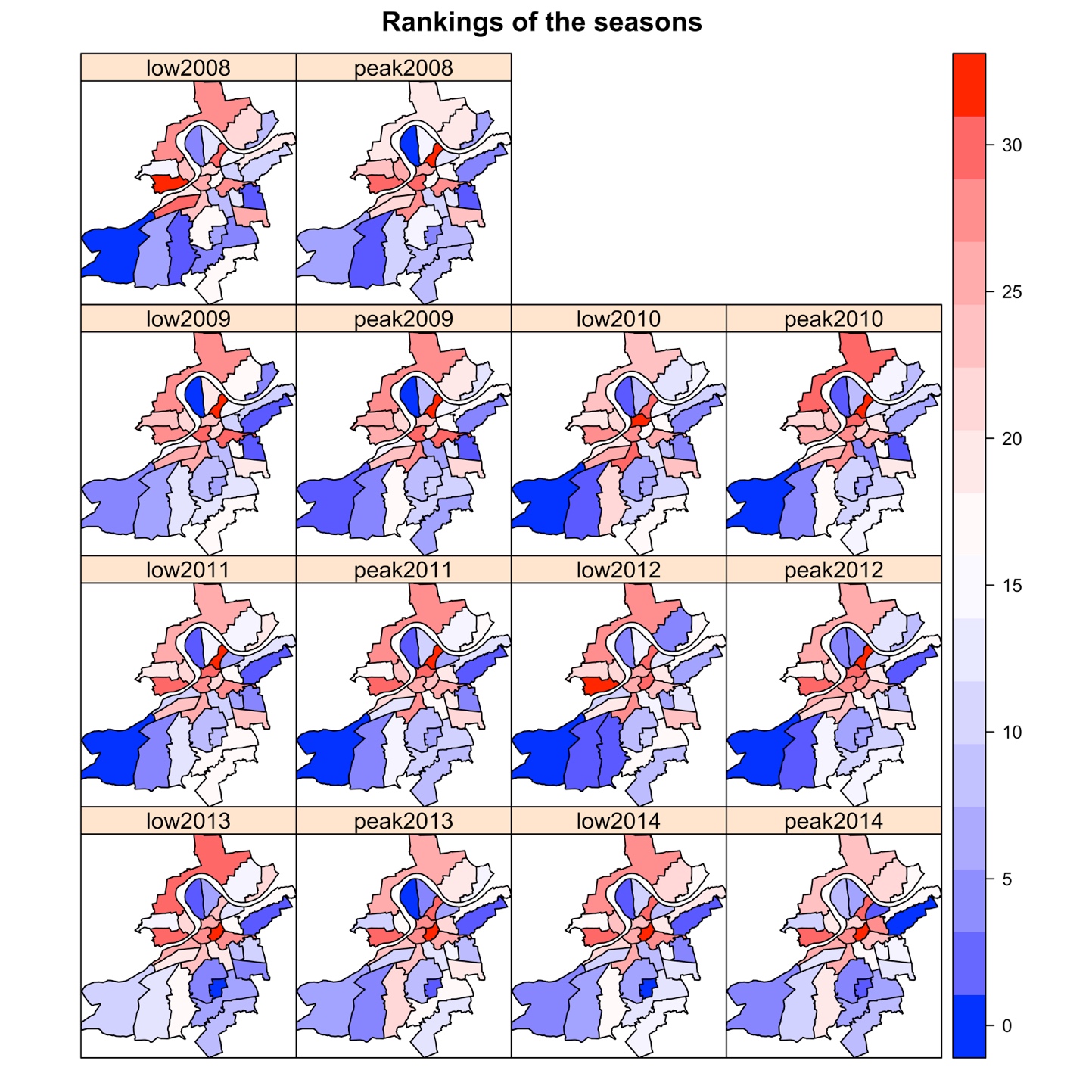

**Fig S1. 7. Spatial patterns malaria.** The panels show the distribution of malaria rankings based on incidence (from low [blue] to high [red] for *P. falciparum*).

**Table S1. 2.** PCA factor loadings contributions of variables (rows) to the different PCs (columns)

| **Variable** | **PC1** | **PC2** | **PC3** |
| --- | --- | --- | --- |
| Total industrial labor | 0.57380 | 19.86171 | 4.52830 |
| Total workers | 2.32322 | 21.77676 | 0.02798 |
| Population density | 1.15284 | 0.33798 | 28.57440 |
| Total Income | 14.50749 | 0.47411 | 7.47029 |
| Household area | 7.83117 | 4.14070 | 1.35731 |
| Water stored | 13.81664 | 1.85728 | 1.05474 |
| Total members | 12.0220 | 1.87349 | 0.00246 |
| Distance | 5.81268 | 0.77895 | 11.58191 |
| Total agriculture labor | 5.15275 | 1.08217 | 7.88543 |
| SC | 1.41002 | 23.06431 | 0.01058 |
| ST | 2.97156 | 20.55982 | 0.08122 |
| Timework | 3.40557 | 1.84227 | 24.78332 |
| Distance to water | 14.38613 | 1.23322 | 2.00411 |
| Scarcity | 14.63408 | 1.11722 | 0.63794 |

**Table S1. 3.** Parameter estimates for the coefficients associated to variables included in the selected model (i.e. explaining more of the variation in the urban malaria cases). The credible intervals (CI) correspond to the 2.5% and 97.5% quantiles of the marginal posterior distribution. Note that the overdispersion parameter of the negative binomial (i.e. the reciprocal of the scale parameter) has a posterior mean value of 2.519 with a 95% credible interval (CI) of [1.456, 3.243]. Thus, the estimated over dispersion parameter (θ) is significantly different from infinity (the value expected for the Poisson special case of the negative binomial).

| ***Covariate*** | **Mean** | **95% CI** |
| --- | --- | --- |
| ***Temperature*** | -0.2001 | [-0.139, -0.307] |
| ***Relative humidity*** | 0.6299 | [0.506, 0.946] |
| ***Principal component 3*** | 0.1055 | [0.039, 0.259] |
| ***Principal component 1*** | 0.546 | [0.369, 0.723] |
| ***Monthly random effect*** | 0.4257 | [0.234, 0.556] |
| ***Yearly random effect*** | 1.2703 | [0.9674, 2.025] |
| ***Spatially-structured hyperparameter*** | 1.926 | [1.681, 2.416] |
| ***Spatially-unstructured hyperparameter*** | 0.926 | [0.681, 1.416] |
| ***Overdispersion parameter*** | 2.519 | [1.4567, 3.243] |

**Table S4.** Predictability evaluation at different levels

| **Model** | **Number of units that predict the same quantile as observed** | **Proportion of units predicting same quantiles as observed** |
| --- | --- | --- |
| Aggregated | 2 | 29% |
| Intermediate | 18 | 57% |
| Disaggregated | 236 | 49% |

**Supplementary text 2**

**Data description and processing**

*Socioeconomic data*

A database of candidate drivers of urban malaria risk in Surat was generated for each reporting unit (zones, units, and workers) for 2010 (the year of the most recent census) using an ordinary kriging method to generate an estimated interpolation surface for each covariate (Supplementary text 2). For this, we used census data from the District Census Handbook at the district level for the year 2010 from the Directorate of Census Operations, Gujarat. We selected variables with a significant effect on malaria transmission, according to the literature [1, 2] (see S1 table 1). In addition, variables such as population density and slum density were calculated using annual population estimates from the Surat Municipal Corporation (SMC). Data on urbanization, housing, water provision, water storage, sanitation, and water scarcity were obtained from water management surveys conducted by Taru Leading Edge, a leading development advisory company in India (<http://taru.co.in/>). This household survey included 90 locations and 400 households (S1 Fig 8), selected to be uniformly distributed throughout the city.

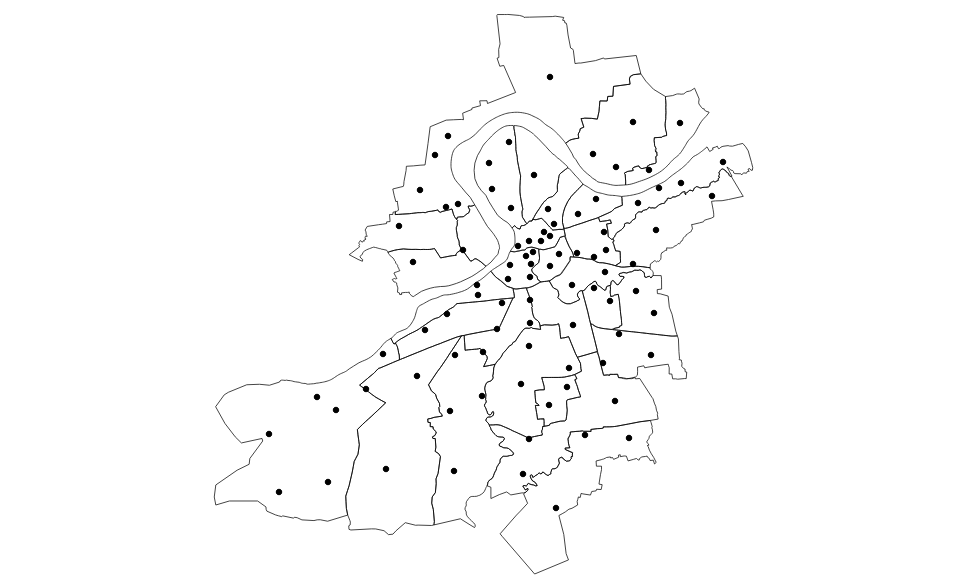

**Figure S1.8.** Spatial distribution of the survey points in the city of Surat.

*Climate data*

We obtained 8-day Land Surface Temperature (LST) from both Terra and Aqua satellites of MODIS for the overlapping period between 2008 and 2015 to estimate surface relative humidity within the city. We used day-time and night-time 8-day composite LST data from MODIS at 1 km spatial resolution (S1 Fig 9). Observed surface air temperature was obtained from the Global Summary of Day [3] dataset of the National Centers for Environmental Information (NCEI), and from 10 meteorological stations located throughout the city for daily temperature, humidity and dew point in the city of Surat since 2014. Although satellite-derived LST offer the advantage of covering a larger area than the local stations, they may experience quality limitations mainly due to cloud contamination. The local station data were therefore used to check for consistency of the satellite products, based only on those stations with continuous long-term data. We calculated the Pearson's correlation coefficient between the interpolated satellite-derived and the meteorological station data, obtaining an R^2^ = 0.72, an indication of high linear correlation between the two data sets at the locations where both are available.

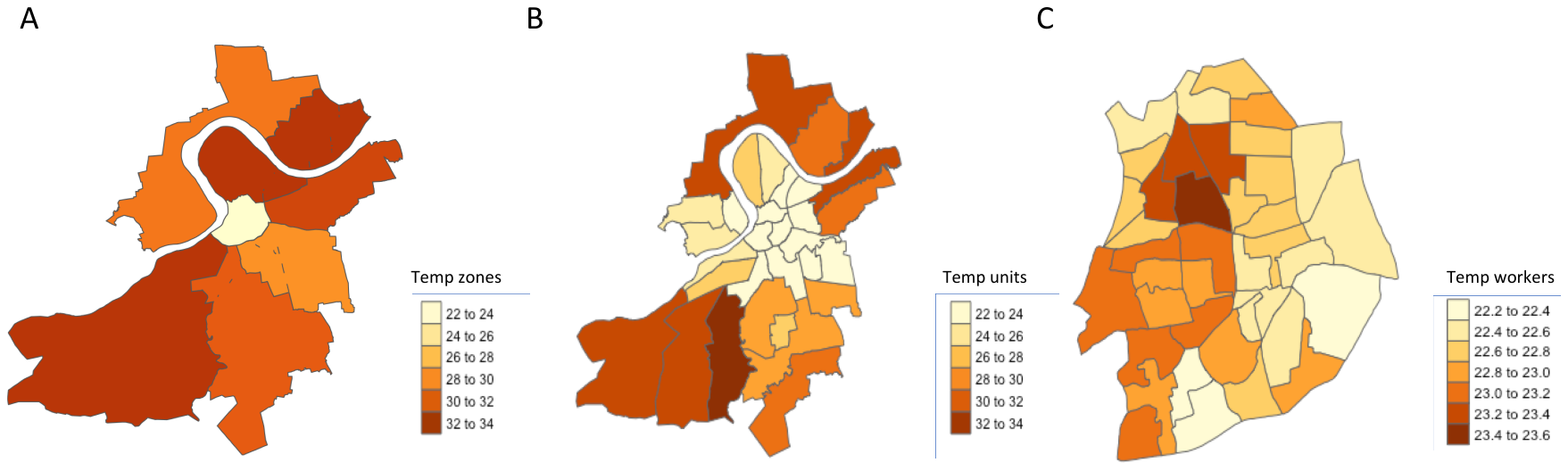

**Figure S1 9.** Variation in the temperature (Land Surface Temperature) extracted from MODIS at different levels of aggregation, A) Zones, B) Units and C) Workers’ Units. For the highest resolution, given the large number of units, only a part of the map is shown here, specifically for the worker units comprising the 3 units at the core of the city.

*Relative humidity generation*

Relative humidity, RH, reflects the amount of moisture in the atmosphere. To estimate surface RH in Surat using MODIS data, meteorological parameters were obtained from ground-based measurements recorded at 10 automatic meteorological stations. MODIS Level-1 data were processed to calculate precipitable water vapor (PW), and MOD07 of MODIS Level-2 atmospheric profile products were used to extract surface air temperature given no cloud cover. These data are derived from known emissivity at different bands of the spectrum, as observed from the satellites in clear sky conditions. The MODIS products also provide associated quality control flags that were used here to screen the quality controlled LST data to further improve the reliability of the datasets.

Data were downloaded from the NASA official website (https://modis.gsfc.nasa.gov/). RH is given by the ratio of vapor pressure (e) and saturation vapor pressure (e_s_) multiplied by 100%. Vapor pressure (e) depends on air pressure (Pa) and specific humidity (Q), while saturation vapor pressure (e_s_) depends on air temperature (Ta) [4]. Their relationship can be described by the following equations:

$RH=\frac{e}{e_{s}}*100$	(1)

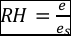

(2)

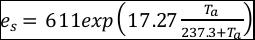

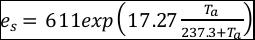

(3)

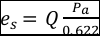

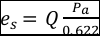

RH can be estimated using precipitable water vapor (PW) retrieved from MODIS. Specific humidity was calculated following the equation:

(4)

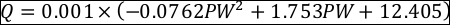

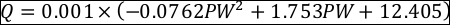

Then air pressure was calculated using:

(5)

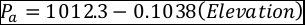

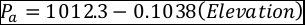

Once specific humidity (Q), air pressure (Pa), air temperature (Ta) are obtained, relative humidity (RH) can be calculated using equations (1)–(3).

**Kriging description**

Kriging is a technique that generates an estimated interpolation surface from a set of data points [5]. This technique incorporates inference from the spatial structure of data points to derive estimations at unmeasured locations. Here we used universal kriging that incorporates spatial trends into the interpolation process. In mathematical terms, the random function Z(x), representing the variable of interest, is split into a deterministic drift m(x) and a random function e’(x) with zero mean

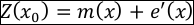

where m(x) is a structural component, associated with a constant mean value or a constant trend and a second-order stationary random function spatially correlated component, known as the variation of the regionalized variable. In universal kriging, the mean is a function of the site coordinates. Then, m(x) follows the equation:

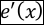

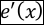

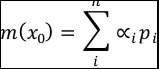

where is the local trend or drift coefficients and is a function of the site coordinates. We assume that the drift is a smooth function of the coordinate vector x and that it can be represented by a linear function where x and y are the coordinates of the *ith* control point and are the drift coefficients.

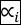

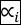

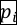

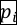

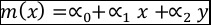

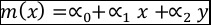

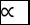

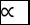

**Interpolation process**

Several forms of kriging interpolators exist: ordinary, universal and simple just to name a few. Here we used an ordinary kriging (OK) interpolation.

a) One assumption that needs to be met in ordinary kriging is that the mean and the variation in the entity being studied is constant across the study area. So, we removed any spatial global trend in our data.
b) For this form of kriging then we calculated the experimental variogram to see the spatial pattern of temperature, relative humidity and socioeconomic data. More specifically, we are interested in how these attribute values (temperature, relative humidity, socioeconomic) vary as the distance between location point pairs increases.
c) Then we fitted a mathematical model to the experimental variogram. Specifically, we were looking for a function that describes the degree of spatial dependence and continuity across data. We generated two theoretical models to best approximate the data. First, a theoretical semi variogram uses a spherical model fit and we fitted an exponential model for the semi variogram.
d) Then we generated a grid of 250mx250m for the Surat area, and with the appropriate variogram model selected we used kriging to predict the respective surfaces for a temperature, relative humidity and socioeconomic variables. The final model uses the measure of variability between points at various distances. Points nearby likely exhibit more similar values, and as the distance between points increases, the similarity between values most likely decreases. In this application, we assume that temperature and socioeconomic measurements that are further apart will vary more than measurements taken close together. We used the Krige function in R to krige the data using the spherical model fit, and plot.

**Supplementary text 3**

**Bayesian modelling**

A negative binomial model was used to allow for overdispersion found in the urban malaria count data [6]. We also incorporated: (i) a spatially unstructured random effect to introduce an extra source of variability/overdispersion in space (a latent effect), and (ii) a spatially structured random effect to explicitly account for spatial autocorrelation and weight relative risk in a region according to the relative risks in neighboring regions. This is consistent with the effect of increased infectious disease risk from neighboring regions of high transmission introduced in both mathematical [7,8,9] and statistical models [10]. We generated two parallel MCMC chains, each of length 25,000 with a burn-in of 20,000 and a thinning of 10, to obtain 1000 samples from the joint posterior distribution. Convergence was assessed by inspecting plots of traces of simulations for individual parameters and monitoring the Gelman-Rubin diagnostic [11]. We standardized the fixed explanatory variables for humidity, temperature, and population density to zero mean and unit variance, to aid MCMC convergence.

Model comparison and evaluation of goodness-of-fit of all the models was assessed at the intermediate level of aggregation (32 units) using the deviance information criterion (DIC) [12], the Watanabe-Akaike information (wAIC) criterion [13], and an statistic for mixed effects models, based on a likelihood ratio (LR) test between the candidate model and an intercept only (null) model [14,15]. Smaller values of DIC and wAIC indicate a better-fitting model. The likelihood ratio R^2^_LR_ ranges from zero to 1, with 1 corresponding to a perfect fit for any reasonable model specification [15]. For each of the resulting models we checked the convergence of the individual parameter estimates and calculated the potential scale reduction. (This reduction is the ratio between the within-chain variance and the posterior variance estimate variances, and values below 1.1 are considered to be acceptable in most cases; see [16] Gelman et al. 2004 for details). New pseudo-observations were simulated by drawing random values from a negative binomial distribution with mean and scale parameter estimated using 10.000 samples from the posterior distribution of the parameters in the model and computing the median cases from the simulations. Finally, model selection was performed at the intermediate spatial resolution (units level, 32 units). We then used this same model formulation at coarser (zone level, 7 units) and finer (worker unit, 486 units) spatial resolutions.

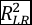

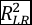
